# Uniparental inheritance promotes adaptive evolution in cytoplasmic genomes

**DOI:** 10.1101/059089

**Authors:** Joshua R. Christie, Madeleine Beekman

## Abstract

Eukaryotes carry numerous asexual cytoplasmic genomes (mitochondria and plastids). Lacking recombination, asexual genomes should theoretically suffer from impaired adaptive evolution. Yet, empirical evidence indicates that cytoplasmic genomes experience higher levels of adaptive evolution than predicted by theory. In this study, we use a computational model to show that the unique biology of cytoplasmic genomes—specifically their organization into host cells and their uniparental (maternal) inheritance—enable them to undergo effective adaptive evolution. Uniparental inheritance of cytoplasmic genomes decreases competition between different beneficial substitutions (clonal interference), promoting the accumulation of beneficial substitutions. Uniparental inheritance also facilitates selection against deleterious cytoplasmic substitutions, slowing Muller’s ratchet. In addition, uniparental inheritance generally reduces genetic hitchhiking of deleterious substitutions during selective sweeps. Overall, uniparental inheritance promotes adaptive evolution by increasing the level of beneficial substitutions relative to deleterious substitutions. When we assume that cytoplasmic genome inheritance is biparental, decreasing the number of genomes transmitted during gametogenesis (bottleneck) aids adaptive evolution. Nevertheless, adaptive evolution is always more efficient when inheritance is uniparental. Our findings explain empirical observations that cytoplasmic genomes—despite their asexual mode of reproduction—can readily undergo adaptive evolution.

## 2 Introduction

About 1.5–2 billion years ago, an *α*-proteobacterium was engulfed by a proto-eukaryote, an event that led to modern mitochondria (Sagan, 1967). Likewise, plastids in plants and algae are derived from a cyanobacterium (Raven and Allen, 2003). These cytoplasmic genomes are essential to extant eukaryotic life, producing much of the energy required by their eukaryotic hosts. Like their ancient ancestors, cytoplasmic genomes reproduce asexually and appear to undergo little recombination with other cytoplasmic genomes (Hagstrom *et al*., 2014; Rokas *et al*., 2003).

Since they lack recombination, asexual genomes have lower rates of adaptive evolution than sexual genomes unless their population size is extremely large (Felsenstein, 1974; Otto and Lenormand, 2002). While the theoretical costs of asexual reproduction have long been known (Felsenstein, 1974; Fisher, 1930; Kondrashov, 1988; Muller, 1932; Otto and Lenormand, 2002), conclusive empirical evidence is more recent (Goddard *et al*., 2005; Lang *et al*., 2013; McDonald *et al*., 2016; Rice and Chippindale, 2001). Three factors largely explain why asexual genomes have low rates of adaptive evolution: (1) beneficial substitutions accumulate slowly; (2) deleterious substitutions are poorly selected against, particularly when their harmful effects are mild; and (3) when beneficial substitutions do spread, any linked deleterious substitutions also increase in frequency through genetic hitchhiking (Felsenstein, 1974; Fisher, 1930; Lang *et al*., 2013; McDonald *et al*., 2016; Muller, 1932).

The lack of recombination in asexual genomes slows the accumulation of beneficial sub-stitutions. Recombination can aid the spread of beneficial substitutions by separating out rare beneficial mutations from deleterious genetic backgrounds (“ruby in the rubbish”) (Peck, 1994). Furthermore, recombination can reduce competition between different beneficial substitutions (“clonal interference”) (Desai and Fisher, 2007; Felsenstein, 1974; Fisher, 1930; Hill and Robertson, 1966; Lang *et al*., 2013; McDonald *et al*., 2016; Muller, 1932; Park and Krug, 2007). Under realistic population sizes and mutation rates, an asexual population will contain multiple genomes—each with different beneficial substitutions—competing with one another for fixation (Desai and Fisher, 2007; Lang *et al*., 2013). Ultimately, clonal interference leads to the loss of some beneficial substitutions, reducing the efficiency of adaptive evolution (Desai and Fisher, 2007; Felsenstein, 1974; Fisher, 1930; Hill and Robertson, 1966; Lang *et al*., 2013; McDonald *et al*., 2016; Muller, 1932; Park and Krug, 2007).

The lack of recombination also makes it more difficult for asexual genomes to purge deleterious substitutions. An asexual genome can only restore a loss of function from a deleterious substitution through a back mutation or a compensatory mutation, both of which are rare (Felsenstein, 1974; Muller, 1964). Unless the size of the population is very large, the number of slightly deleterious substitutions should increase over time as the least-mutated class of genome is lost through genetic drift (‘Muller’s ratchet”) Felsenstein, 1974; Muller, 1964).

If that were not enough, asexual genomes are also especially susceptible to genetic hitch-hiking (Lang *et al*., 2013; McDonald *et al*., 2016), a process by which deleterious sub-stitutions spread through their association with beneficial substitutions (Gillespie, 2000; Smith and Haigh, 1974). As all loci on an asexual genome are linked, deleterious and beneficial substitutions on the same genome will segregate together. When the positive effect of a beneficial substitution outweighs the negative effect of a deleterious substitution, the genome that carries both can spread through positive selection (Gillespie, 2000; Smith and Haigh, 1974). Even when the additive effect is zero or negative, a beneficial substitution can still aid the spread of a deleterious substitution via genetic drift by reducing the efficiency of selection against the deleterious substitution. Genetic hitch-hiking can thus offset the benefits of accumulating beneficial substitutions by interfering with the genome’s ability to purge deleterious substitutions (Gillespie, 2000; Smith and Haigh, 1974).

Free-living asexual organisms generally have very large population sizes (Mamirova *et al*., 2007) and may undergo occasional sexual exchange (e.g. conjugation in bacteria (Narra and Ochman, 2006)), allowing these organisms to alleviate some of the costs of asexual reproduction (Felsenstein, 1974; Otto and Lenormand, 2002). Asexual cytoplasmic genomes, however, have an effective population size much smaller than that of free-living asexual organisms (Ballard and Whitlock, 2004; Mamirova *et al*., 2007). As a smaller population size increases the effect of genetic drift, cytoplasmic genomes should have less efficient selection than asexual organisms (Lynch *et al*., 2006; Neiman and Taylor, 2009) and should struggle to accumulate beneficial substitutions and to purge deleterious substitutions (Birky, 2008; Lynch, 1996; Rispe and Moran, 2000).

Although there are indications that cytoplasmic genomes suffer from these costs of asexual reproduction (e.g. low binding stability of mitochondrial transfer RNAs (Lynch, 1996)), cytoplasmic genomes also readily undergo adaptive evolution, particularly in animals. Animal mitochondrial protein-coding genes show signatures that are consistent with both low levels of deleterious substitutions (Cooper *et al*., 2015; Mamirova *et al*., 2007; Popadin *et al*., 2013) and frequent selective sweeps of beneficial substitutions (Bazin *et al*., 2006; Meiklejohn *et al*., 2007). Indeed, it is estimated that 26% of mitochondrial substitutions that alter proteins in animals have become fixed through adaptive evolution (James *et al*., 2016). Beneficial substitutions in the mitochondrial genome have helped animals adapt to specialized metabolic requirements (Castoe *et al*., 2008; da Fonseca *et al*., 2008; Grossman *et al*., 2004; Shen *et al*., 2010) and have enabled humans to adapt to cold northern climates (Ruiz-Pesini *et al*., 2004). Likewise, it is clear that adaptive evolution has played a role in the evolution of plastid genomes (Cui *et al*., 2006; Zhong *et al*., 2009).

How then do we reconcile empirical evidence for adaptive evolution in cytoplasmic genomes with theoretical predictions that such adaptation should be impaired? Un-like free-living asexual organisms, which are directly exposed to selection, cytoplasmic genomes exist within host cells. The fitness of cytoplasmic genomes is therefore closely aligned with the fitness of their host. Each of these hosts carries multiple cytoplasmic genomes that are generally inherited from a single parent (uniparental inheritance) (Christie *et al*., 2015). During gametogenesis, cytoplasmic genomes can undergo tight population bottlenecks, affecting the transmission of genomes from parent to offspring (Birky, 1995; Cao *et al*., 2007). Cytoplasmic genomes are thus subject to very different evolutionary pressures than free-living asexual organisms.

Some of the effects of uniparental inheritance and a transmission bottleneck on the evolution of cytoplasmic genomes have already been identified. Both uniparental inheritance and a transmission bottleneck decrease within-cell variance in cytoplasmic genomes and increase between-cell variance. (Bergstrom and Pritchard, 1998; Christie *et al*., 2015; Hadjivasiliou *et al*., 2013; Roze *et al*., 2005). Uniparental inheritance is known to select against deleterious mutations (Hadjivasiliou *et al*., 2013; Hastings, 1992; Roze *et al*., 2005) and select for mito-nuclear coadaptation (Hadjivasiliou *et al*., 2012). Similarly, a transmission bottleneck and other forms of within-generation drift are known to slow the accumulation of deleterious substitutions in cytoplasmic genomes (Bergstrom and Pritchard, 1998; Rispe and Moran, 2000; Takahata and Slatkin, 1983).

Although the effect of uniparental inheritance and a bottleneck on the accumulation of deleterious substitutions is reasonably well-studied, much less attention has been paid to the other limitations of asexual reproduction: slow accumulation of beneficial sub-stitutions and high levels of genetic hitchhiking. The two studies that have addressed the spread of beneficial substitutions have come to contradictory conclusions. Takahata and Slatkin (Takahata and Slatkin, 1983) showed that within-generation drift promoted the accumulation of beneficial substitutions. In contrast, Roze and colleagues (Roze *et al*., 2005) found that within-generation drift due to a bottleneck reduced the fixation probability of a beneficial mutation. Takahata and Slatkin found no difference between uniparental and biparental inheritance of cytoplasmic genomes (Takahata and Slatkin, 1983) while Roze and colleagues found that uniparental inheritance increased the fixation probability of a beneficial mutation and its frequency at mutation-selection equilibrium (Roze *et al*., 2005). Of the two previous studies, only the model of Takahata and Slatkin was able to examine the accumulation of substitutions (Takahata and Slatkin, 1983) (the model of Roze and colleagues only considered a single locus (Roze *et al*., 2005)). To our knowledge, no study has looked at how inheritance mode affects genetic hitchhiking in cytoplasmic genomes.

Here we develop theory that explains how cytoplasmic genomes are capable of adaptive evolution despite their lack of recombination. We will show how the biology of cytoplasmic genomes—specifically their organization into host cells and their uniparental inheritance—can allow them to accumulate beneficial substitutions and to purge deleterious substitutions very efficiently compared to free-living asexual genomes.

## 3 Model

For simplicity, we base our model on a population of diploid single-celled eukaryotes. We examine the accumulation of beneficial and deleterious substitutions in an individual-based computational model that compares uniparental inheritance of cytoplasmic genomes with biparental inheritance. Since we are interested in the evolutionary consequences of each trait, rather than the evolution of the traits, we examine each form of inheritance separately. As genetic drift plays an important role in the spread of substitutions, we take stochastic effects into account. We vary the size of the transmission bottleneck during gametogenesis (i.e. the number of cytoplasmic genomes passed from parent to gamete) to alter the level of genetic drift. To examine how the organization of cytoplasmic genomes into host cells affects their evolution, we also include a model of comparable free-living asexual genomes.

We have four specific aims. We will determine how inheritance mode and the size of the transmission bottleneck affect (Aim 1) clonal interference and the accumulation of beneficial substitutions; (Aim 2) the accumulation of deleterious substitutions; (Aim 3) the level of genetic hitchhiking; and (Aim 4) the level of adaptive evolution, which we define as the ratio of beneficial to deleterious substitutions. Although uniparental inheritance and a transmission bottleneck are known to select against deleterious mutations on their own (Bergstrom and Pritchard, 1998; Hadjivasiliou *et al*., 2013; Hastings, 1992; Roze *et al*., 2005; Takahata and Slatkin, 1983), the interaction between inheritance mode, transmission bottleneck, and the accumulation of deleterious substitutions has not to our knowledge been examined. Thus we include Aim 2 to specifically examine interactions between inheritance mode and size of the transmission bottleneck. To address our aims, we built four variations of our model. First, we examine clonal interference and the accumulation of beneficial substitutions using a model that considers beneficial but not deleterious mutations (Aim 1). Second, we consider deleterious but not beneficial mutations to determine how inheritance mode and a transmission bottleneck affect the accumulation of deleterious substitutions in cytoplasmic genomes (Aim 2). Third, we combine both beneficial and deleterious substitutions. This allows us to examine the accumulation of deleterious substitutions in the presence of beneficial mutations (genetic hitchhiking; Aim 3) and the ratio of beneficial to deleterious substitutions (Aim 4). For all aims, we compare our models of cytoplasmic genomes to a comparable population of free-living asexual genomes. This serves as a null model, allowing us to examine the strength of selection when asexual genomes are directly exposed to selection.

### 3.1 Cytoplasmic genome model

The population contains N individuals, each carrying the nuclear genotype *Aa*, where *A* and *a* are self-incompatible mating type alleles. Diploid cells contain *n* cytoplasmic genomes, and each genome has *l* linked base pairs. A cytoplasmic genome is identified by the number of beneficial and deleterious substitutions it carries (α and κ respectively; note, we do not track where on the genome the mutations occur). Cells are identified by the number of each type of cytoplasmic genome they carry. The life cycle has four stages, and a complete passage through the four stages represents a generation. The first stage is **mutation**. Initially, all cells carry cytoplasmic genomes with zero substitutions. Mutations can occur at any of the *l* base pairs. The probability that one of these l sites will mutate to a beneficial or deleterious site is given by *μ*_b_ and *μ*_d_ per site per generation respectively (determined via generation of random numbers within each simulation).

After mutation, cells are subject to **selection**, assumed for simplicity to act only on diploid cells. We assume that each substitution has the same effect, which is given by the selection coefficient (*s*_*b*_ for beneficial and *s*_*d*_ for deleterious) and that fitness is additive. We assume that a cell’s fitness depends solely on the total number of substitutions carried by its cytoplasmic genomes. Cells are assigned a relative fitness based on the number of beneficial and deleterious substitutions carried by their cytoplasmic genomes. These fitness values are used to sample N new individuals for the next generation.

Each of the post-selection diploid cells then undergoes **gametogenesis** to produce two gametes, one with nuclear allele *A* and the other with nuclear allele *a*. Each gamete also carries *b* cytoplasmic genomes sampled with replacement from the *n* cytoplasmic genomes carried by the parent cell (with *b* ≤ *n*/2). We examine both a tight transmission bottleneck (few genomes are transmitted) and a relaxed transmission bottleneck (more genomes are transmitted). To maintain the population size at *N*, each diploid cell produces two gametes.

During **mating**, each gamete produced during gametogenesis is randomly paired with another gamete of a compatible mating type. These paired cells fuse to produce diploid cells. Under biparental inheritance, both the gametes with the *A* and a alleles pass on their *b* cytoplasmic genomes, while under uniparental inheritance, only the *b* genomes from the gamete with the *A* allele are transmitted. Finally, *n* genomes are restored to each new diploid cell by sampling n genomes with replacement from the genomes carried by the diploid cell after mating (2*b* under biparental inheritance and b under uniparental inheritance). The model then repeats, following the cycle of mutation, selection, game-togenesis, and mating described above.

### 3.2 Free-living genome model

To clarify how the organization of cytoplasmic genomes into hosts affects their evolution, we also examine a model of free-living asexual cells. We examine two different population sizes for free-living cells: (1) *N*_*FL*_ = *N* × *n* (matched to the number of cytoplasmic genomes); or (2) *N*_*FL*_ = *N* (matched to the number of eukaryotic hosts). Each free-living cell carries one haploid asexual nuclear genome with l base pairs. Now there are only two stages to the life cycle: mutation and selection. Mutation proceeds as in the model of cytoplasmic genomes. Selection, however, now depends only on the number of substitutions carried by the single free-living genome.

As the fitness effect of a mutation in a free-living cell’s genome is not directly comparable to the fitness effect of a mutation in a host’s cytoplasmic genomes, we examine a range of possibilities. As a default, we assume that each mutation in a free-living cell’s genome impacts its fitness by the same magnitude as each mutation on a cytoplasmic genome impacts its host’s fitness (e.g. the fitness of a free-living cell that carries a single beneficial substitution is equivalent to the fitness of a host that carries a single beneficial substitution on one of its cytoplasmic genomes). However, since cytoplasmic genomes exist in multiple copies within a host, a single substitution on a single cytoplasmic genome might impact fitness less than a single substitution on a free-living genome (Haig, 2016). To address this, we vary the effect of substitutions on fitness in free-living genomes relative to cytoplasmic genomes. The parameter s_F L_ represents the effect of substitutions on free-living fitness relative to cytoplasmic genomes (e.g. *s*_*FL*_ = 10 means that a single substitution in a free-living genome has a 10-fold greater effect on free-living fitness than a single substitution on a single cytoplasmic genome has on host fitness). Our intention is not to accurately model extant populations of free-living asexual organisms, as these differ in a number of ways from cytoplasmic genomes (e.g. population size, mutation rate, and genome size (Mamirova *et al*., 2007)), but rather to examine how the organization of multiple cytoplasmic genomes within a host affects their evolution.

### 3.3 Parameter value estimates

Our default population size is *N* = 1000, number of mitochondria is *n* = 50, and size of the transmission bottleneck is either *b* = *n*/2 (relaxed bottleneck) or *b* = *n*/10 (tight bottleneck). A value of *n* = 50 is frequently used in models of mitochondrial evolution (Christie *et al*., 2015; Hadjivasiliou *et al*., 2012, 2013; Hastings, 1992). When *n* = 50 and either a tight or relaxed bottleneck is applied, the number of resulting cytoplasmic genomes (5–25) corresponds to the number of mitochondria or plastids in the gametes of isogamous species such as *Physarum polycephalum* (Moriyama and Kawano, 2003), *Saccharomyces cerevisiae* (Hoffmann and Avers, 1973), and *Chlamydomonas reinhardtii* (Nishimura *et al*., 1998). We also examine *n* = 200, which results in a transmission bottleneck size similar to that in animals (Jenuth *et al*., 1996; Wai *et al*., 2008).

We fix the number of base pairs at *l* = 20; 000, which is roughly the size of the animal mitochondrial genome (Boore, 1999). As the mutation rate in animal mitochondrial DNA (mtDNA) is between 7:8×10^−8^ and 1.7×10^−7^ per nucleotide per generation (Denver *et al*., 2000; Haag-Liautard *et al*., 2008; Xu *et al*., 2012), we let *μ*_*d*_ = 1×10^−7^ per nucleotide per generation, under the assumption that the majority of mutations are deleterious (Eyre-Walker and Keightley, 2007). Although we are not aware of any direct estimates for the rate of beneficial mutations in mitochondrial DNA, studies have estimated the relative proportion of mutations that are beneficial in other types of genomes. These beneficial mutation estimates range from undetectable (in the bacteriophage *φ*6 (Burch *et al*., 2007), the yeast *Saccharomyces paradoxus* (Koufopanou *et al*., 2015), and *Escherichia coli* (Elena *et al*., 1998)), to moderately common (6% in Saccharomyces cerevisiae (Joseph and Hall, 2004), 4% in the vesicular stomatitis virus (Sanjuán *et al*., 2004), 15% in the bacteriophage *φ*X174 (Silander *et al*., 2007)), to extremely common (25% of fitness-altering mutations in *Saccharomyces cerevisiae* (Dickinson, 2008) and ≈50% of fitness-altering mutations in *Arabidopsis thaliana* (Shaw *et al*., 2000)). We examine beneficial mutations that are rare (*μ*_*b*_ = 1×10^−9^ per nucleotide per generation; 1% of the deleterious mutation rate) to moderately common (*μ*_*b*_ = 1×10^−8^ per nucleotide per generation; 10% of the deleterious mutation rate).

We focus on selection coefficients that represent mutations with small effects on fitness: *s*_*b*_ = 0:01−0:1 (see the legend of Figure 1 for a description of how the selection coefficient translates to individual fitness). Since it is difficult to estimate the relative impact on fitness of a mutation on a free-living genome compared to mutation on a cytoplasmic genome, we let *s*_*FL*_ vary from 1–50.

**Figure 1:**
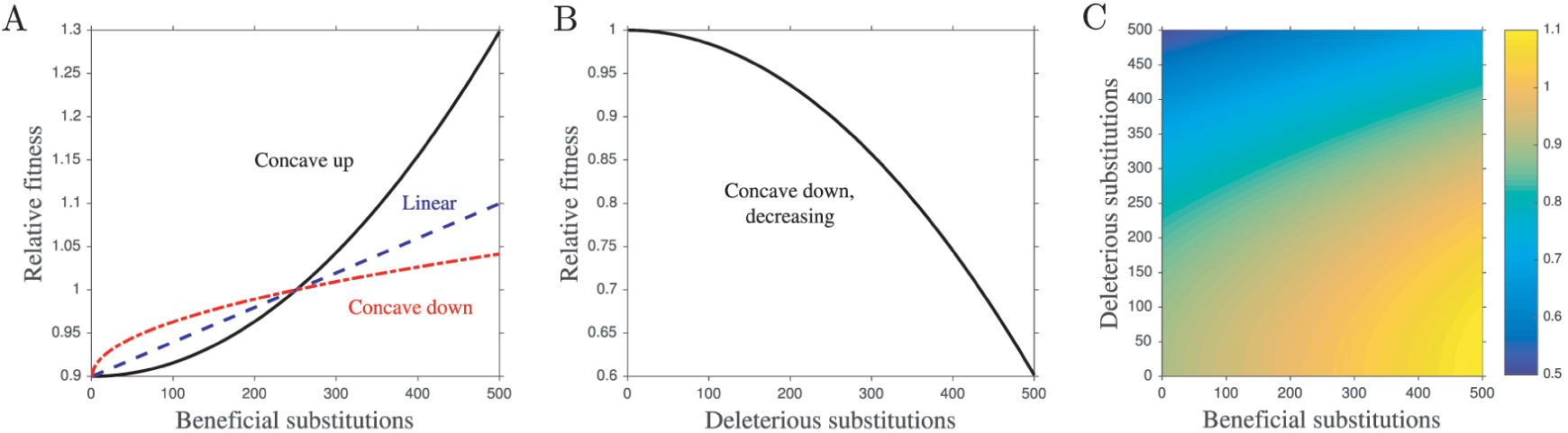
Fitness functions. Additional parameters: *n* = 50, *s*_b_ = 0:1, *s*_*d*_ = 0:1, γ = 5. **A**. The three fitness functions used in this study in the case of beneficial mutations only. The selection coefficient is defined such that 1−*s*_*b*_ represents the fitness of a cell with zero beneficial substitutions (a cell with *nγ* beneficial substitutions has a fitness of 1, where *n* is the number of cytoplasmic genomes and *γ* is the number of substitutions each cytoplasmic genome must accumulate before the simulation is terminated). In this example, where *n* = 50, *s*_*b*_ = 0:1, and *γ* = 5, a cell’s fitness is 0.9 when its cytoplasmic genomes carry no beneficial substitutions, and its fitness is 1 when each cytoplasmic genome in the cell carries an average of 5 substitutions (50 × 5 = 250 beneficial substitutions in total). **B**. The deleterious fitness function. Here, a cell with no deleterious substitutions has a fitness of 1, while a cell with *nγ* substitutions has a fitness of 1−*s*_*d*_. We only examine a concave down decreasing function for the accumulation of deleterious substitutions (unless we are comparing cytoplasmic genomes to free-living genomes, in which case we use a linear fitness function). **C**. One of the fitness functions used in the model with both beneficial and deleterious mutations. The beneficial substitution portion of the function can take any of the forms in panel **A** while the deleterious substitution portion takes the form in panel **B** (unless we are comparing cytoplasmic genomes to free-living genomes, in which case both the beneficial and deleterious fitness functions are linear). In this example the fitness surface combines a linear function for beneficial substitutions with a concave down fitness function for deleterious substitutions. The color represents the fitness of a cell carrying a given number of deleterious substitutions (x-axis) and beneficial substitutions (y-axis). Equations for the fitness functions can be found in section S3.2 (**A**), section S4 (**B**), and section S5.2. (**C**).

As there are few data on the distribution of fitness effects of beneficial substitutions in cytoplasmic genomes, we examine three fitness functions: concave up, linear, and concave down (Figure 1A). For deleterious substitutions in cytoplasmic genomes, there is strong evidence that fitness is only strongly affected when the cell carries a high proportion of deleterious genomes (Chinnery and Samuels, 1999; Rossignol *et al*., 2003), and so we use a decreasing concave down function to model deleterious substitutions (Figure 1B). When we combine beneficial and deleterious mutations in a single model, we examine the three fitness functions for the accumulation of beneficial substitutions but only a concave down decreasing fitness function for the accumulation of deleterious substitutions (Figure 1B). When comparing free-living and cytoplasmic genomes, we always use a linear fitness function for both beneficial and deleterious substitutions because for this function the strength of selection on a new substitution is independent of existing substitution load.

In the model that considers beneficial mutations only (Aim 1), the simulation stops once every cytoplasmic genome in the population has accumulated at least γ beneficial substitutions. For the remaining models, each simulation runs for 10,000 generations. For all models, we average the results of 500 Monte Carlo simulations for each combination of parameter values (we vary *N*, *n*, *b*, *s*_*b*_, *s*_*d*_, *s*_*FL*_, and the fitness functions associated with beneficial substitutions). We wrote our model in R version 3.1.2 (Team, 2013). For a detailed description of the models, see section S3—section S5.

## 4 Results

### 4.1 Uniparental inheritance of cytoplasmic genomes promotes the accumulation of beneficial substitutions

For conceptual purposes, we break down the accumulation of beneficial substitutions into two phases. We call the first the “drift phase”. In this phase, the genome type with α substitutions continuously arises in a population that contains genomes with α − 1 or fewer beneficial substitutions, but it is repeatedly lost to drift and does not spread (since we examine small selection coefficients, drift dominates the fate of genomes when they are rare). The drift phase starts when we first observe a genome with α substitutions and ends when that genome persists in the population (i.e. it is no longer lost to drift).

The second phase, which we call the “selection phase”, involves the spread of the genome with α substitutions through positive selection. The selection phase commences at the end of the drift phase (i.e. once the genome with *α* substitutions persists in the population) and ends when a genome carrying *α*+1 substitutions first appears in the population. At this point, the drift phase of the genome with *α*+1 substitutions begins and the cycle continues.

Gametogenesis introduces variation in the cytoplasmic genomes that are passed to gametes. Gametes can thus carry a higher or lower proportion of beneficial substitutions than their parent. Uniparental inheritance maintains this variation in offspring, reducing within-cell variation (Figure 2A) while increasing between-cell variation (Figure 2B). Biparental inheritance, however, combines the cytoplasmic genomes of different gametes, destroying much of the variation produced during gametogenesis and reducing between-cell variation (Figure 2B). Thus, selection is more efficient when inheritance is uniparental because there is more between-cell variation in fitness on which selection can act (Figure 2B).

**Figure 2:**
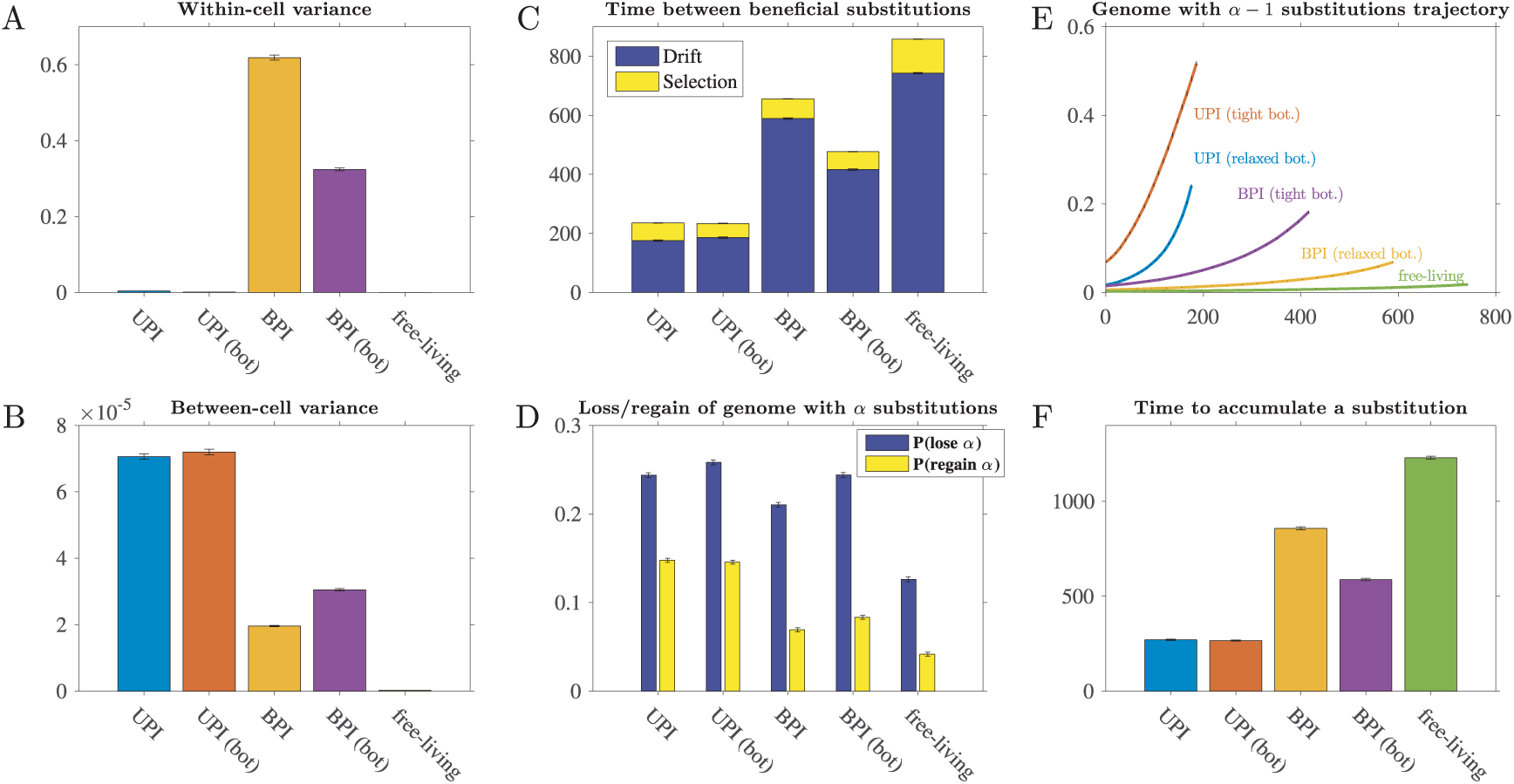
Dynamics in the accumulation of beneficial substitutions. Parameters: *N* = 1000, *n* = 50, *s*_*b*_ = 0:1, *μ*_b_ = 10^−8^, linear fitness function, and *b* = 25 (relaxed transmission bottleneck) or *b* = 5 (tight transmission bottleneck). Error bars represent standard error of the mean. UPI: uniparental inheritance with a relaxed bottleneck, UPI (bot): uniparental inheritance with a tight bottleneck, BPI: biparental inheritance with a relaxed bottleneck, and BPI (bot): biparental inheritance with a tight bottleneck. **A**. Variance in the number of different cytoplasmic genomes carried by cells (averaged over all cells in the population each generation). As free-living cells carry a single genome, they have no within-cell variance. **B**. Variance of all cells’ fitness values (averaged over each generation). (Note that between-cell variation in the free-living population is depicted but is so low that it appears as zero.) **C**. The number of generations separating the genome carrying α substitutions from the genome carrying α + 1 (averaged over all observed substitutions, but excluding α = 1, as the dynamics of α = 1 are largely driven by the starting conditions). In the drift phase, depicted in dark blue, the genome carrying α substitutions arises but is lost to drift. In the selection phase, depicted in yellow, the genome with α substitutions spreads through positive selection (see main text for a detailed description of the drift and selection phases). During the drift phase of the genome with α substitutions, **D** shows the probability of losing all genomes with α substitutions (*P*(lose *α*)) and the probability of regenerating at least one genome with *α* substitutions once all genomes with *α* substitutions have been lost (*P*(regain *α*)) (averaged over all observed drift periods, but excluding *α* = 1). During the drift phase of the genome with *α* substitutions, **E** shows the trajectory of the genome with *α* − 1 substitutions. To calculate the curves, we divided each of the 500 Monte Carlo simulations into 20 equidistant pieces. We rounded to the nearest generation and obtained the frequency of the genome with *α* − 1 substitutions at each of those 20 generation markers. Each curve shows the average of those 20 generation markers (over all drift phases, excluding *α* = 1, and over all simulations) and is plotted so that the end of the curve aligns with the mean length of the drift phase (shown in panel **C**). **F**. The mean number of generations to accumulate a single beneficial substitution (*s*_*FL*_ = 1 for free-living). We divide the number of generations to accumulate *γ* substitutions by the mean number of beneficial substitutions accumulated in that time period (averaged over all simulations).

Under uniparental inheritance, it takes less time for the genome with *α* substitutions to generate the genome with *α* + 1 substitutions than under biparental inheritance (Figure 2C). Uniparental inheritance reduces the time that the genome with *α* substitutions spends in the drift phase (Figure 2C) by increasing the rate at which the genome with *α* substitutions is regenerated once lost to drift (Figure 2D). The regeneration of the genome with α substitutions is proportional to the rate at which mutations occur on the genome with *α* − 1 substitutions, which in turn is proportional to the frequency of the genome with *α* − 1 substitutions in the population. Under uniparental inheritance, the genome with *α* − 1 substitutions increases in frequency much more quickly than under biparental inheritance (Figure 2E), presenting a larger target for de novo mutations and driving regeneration of the genome with *α* substitutions (Figure 2D). As a result, under uniparental inheritance cytoplasmic genomes suffer less from clonal interference (Figure 3) and take less time to accumulate beneficial substitutions than under biparental inheritance (Figure 2F; see Figure S1 for a range of different parameter values).

**Figure 3:**
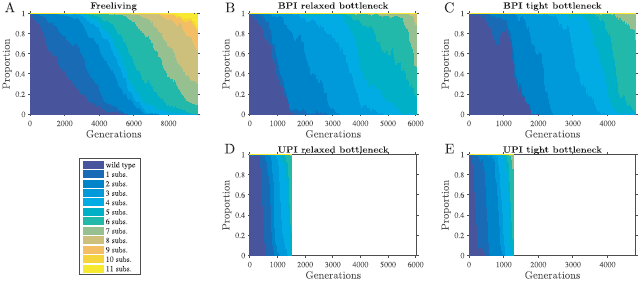
Uniparental inheritance reduces clonal interference. Parameters: *N* = 1000, *n* = 50, *s*_*b*_ = 0:1, *μ*_*b*_ = 10^−8^, and a linear fitness function. The figure depicts a time-series of a single simulation, showing the proportions of genomes carrying different numbers of substitutions (we chose the first completed simulation for each comparison). We report a linear approximation of the mean slope of declines in proportion of the wild type genome as *m*_*g*_. (*m*_*g*_ has units of %/generation and is determined by dividing −99:5% by the mean number of generation for the wild type genome to drop from 100% to below 0.5%.) We also report the mean number of genomes co-existing in the population, which we call *c*_*g*_. **A**. In a population of free-living cells, genomes with beneficial substitutions spread slowly through the population (*m*_*g*_ = −0:017 %=generation). As a result, multiple genomes co-exist at any one time (*c*_*g*_ = 7:0 genomes), increasing the scope for clonal interference. **B–C**. Biparental inheritance with a relaxed bottleneck (**B**; *b* = 25) and tight bottleneck (**C**; *b* = 5). Under biparental inheritance, genomes carrying beneficial substitutions spread more quickly compared to free-living genomes (**B**: *m*_*g*_ = −0:039 %=generation; **C**: *m*_*g*_ = −0:072 %=generation), reducing the number of co-existing genomes (**B**: *c*_*g*_ = 4:8 genomes; **C**: *c*_*g*_ = 3:8 genomes). **D–E**. Uniparental inheritance with a relaxed bottleneck (**D**; *b* = 25) and tight bottleneck (**E**; *b* = 5). Under uniparental inheritance, genomes with beneficial substitutions spread much more quickly than free-living and biparentally inherited cytoplasmic genomes (**D**: *m*_*g*_ = −0:215 %=generation; **E**: *m*_*g*_ = −0:220 %=generation). This leads to fewer genomes co-existing in the population (**D**: *c*_*g*_ = 3:1 genomes; E: cg = 2:8 genomes) and low levels of clonal interference.

### 4.2 Cytoplasmic genomes generally accumulate beneficial mutations faster than free-living genomes

The units of selection differ between cytoplasmic genomes (eukaryotic host cell) and free-living genomes (free-living asexual cell). Cytoplasmic genomes have two levels at which variance in fitness can be generated: variation in the number of substitutions per genome and variation in the relative number of each genome type in a host cell (Figure 2A). In contrast, free-living genomes can differ only in the number of substitutions carried per genome. Consequently, when a mutation on a cytoplasmic genome has the same effect as a mutation on a free-living genome (i.e. *s*_*FL*_ = 1), cytoplasmic genomes have a greater potential for creating variance between the units of selection than free-living genomes (Figure 2B).

In cytoplasmic genomes, the genome with α substitutions spends less time in the drift phase compared to free-living genomes when *s*_*FL*_ = 1 (Figure 2C). Cytoplasmic genomes have a shorter drift phase not because they are less likely to be lost by drift—in fact cytoplasmic genomes are more frequently lost to drift than free-living genomes—but because once a genome with α substitutions has been lost, it is more quickly regenerated (Figure 2D). Since cytoplasmic genomes experience strong positive selection (Figure 2B), cytoplasmic genomes with *α* − 1 substitutions quickly increase in frequency (Figure 2E), driving the formation of the genome with *α* substitutions. As a result, cytoplasmic genomes have lower levels of clonal interference (Figure 3), reducing the time to accumulate beneficial substitutions compared to free-living genomes when *s*_*FL*_ = 1 (Figure 2F).

When mutations on a free-living genome have a larger effect on fitness compared to mutations on a cytoplasmic genome (i.e. *s*_*FL*_ > 1), free-living genomes can accumulate beneficial substitutions more quickly than cytoplasmic genomes with uniparental inheritance (Figure 4). When we match the population size of free-living genomes to the number of eukaryotic hosts, free-living genomes accumulate beneficial substitutions at a lower rate than cytoplasmic genomes unless mutations in free-living genomes have a 50-fold effect on fitness (Figure 4A). When we match the population size of free-living genomes to the number of cytoplasmic genomes, free-living genomes accumulate beneficial substitutions more quickly than cytoplasmic genomes when mutations in free-living genomes have a 20-fold or greater effect on fitness (Figure 4B). Beneficial substitutions accumulate more quickly in larger populations of free-living genomes (Figure 4); in larger populations, beneficial mutations arise more frequently and are less susceptible to genetic drift.

**Figure 4:**
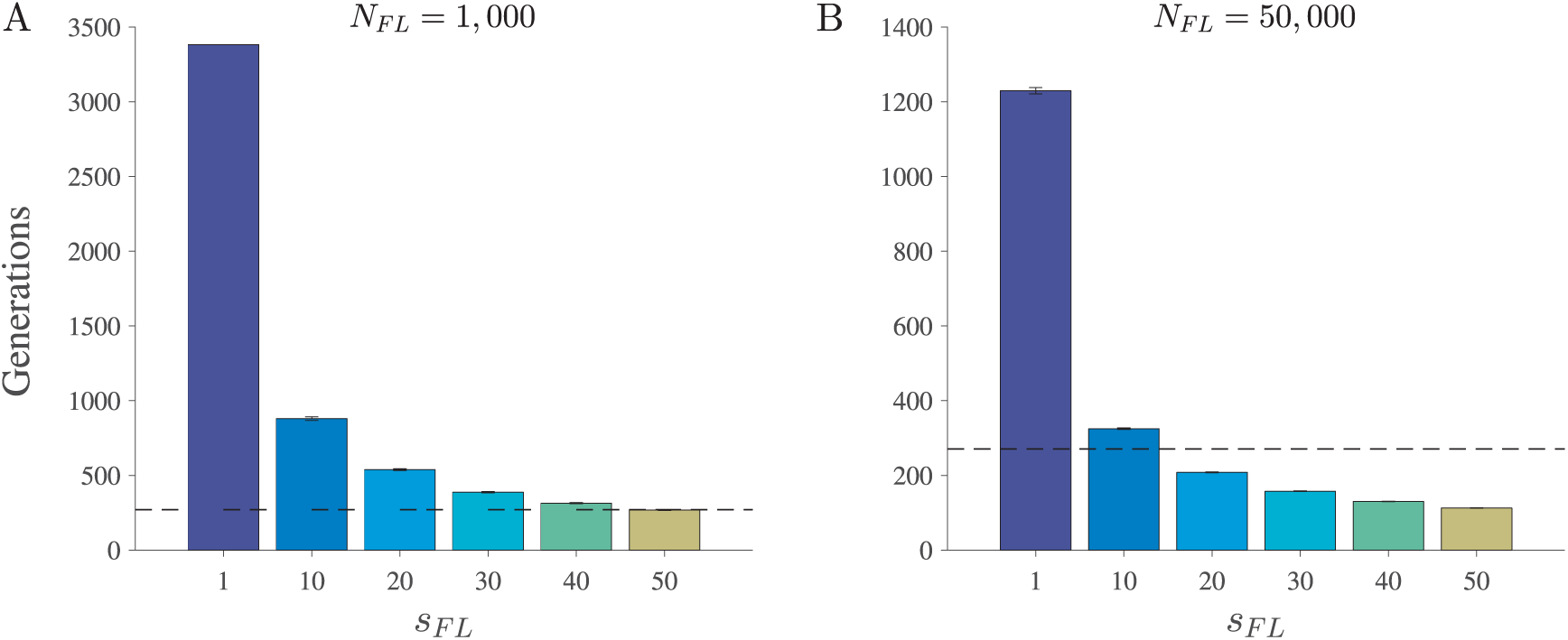
Varying the effect of beneficial substitutions on fitness of free-living cells. Parameters: *s*_*b*_ = 0:1, *γ* = 5, μ_*b*_ = 10^−8^ and a linear fitness function. When *N*_*FL*_ = 1000, the population size of free-living genomes is equal to the number of eukaryotic hosts; when *N*_*FL*_ = 50; 000, the population size of free-living genomes is equal to the number of cytoplasmic genomes (assuming *N* = 1000 and *n* = 50, as in Figure 2). The y-axis shows the mean number of generations to accumulate a single beneficial substitution (see Figure 2F legend for details). On the x-axis, we vary the effect mutations have on the fitness of free-living cells. A mutation on a free-living genome has an *s*_*FL*_-fold effect on its cell’s fitness compared to the effect of a mutation on a cytoplasmic genome on its host’s fitness. The dashed line represents the mean number of generations required to accumulate a beneficial substitution assuming uniparental inheritance (relaxed bottleneck) under equivalent conditions (≈ 272; see Figure 2F). **A**. Population size of free-living genomes equals 1000. **B**. Population size of free-living genomes equals 50,000. Error bars are ± standard error of the mean.

### 4.3 Inheritance mode is more important than the size of the bottleneck

Under biparental inheritance, a tight bottleneck decreases the variation in cytoplasmic genomes within gametes (Figure 2A) and increases the variation between gametes (Figure 2B). Consequently, under biparental inheritance beneficial substitutions accumulate more quickly than when the transmission bottleneck is relaxed (Figure 2F and Figure S1). Bottleneck size has less of an effect on uniparental inheritance because uniparental inheritance efficiently maintains the variation generated during gametogenesis even when the bottleneck is relaxed (Figure 2B). When *n* is larger (*n* = 200), a tight bottleneck reduces the time for beneficial substitutions to accumulate, but even here the effect is minor (Figure S1C).

Importantly, the accumulation of beneficial substitutions under biparental inheritance and a tight bottleneck is always less effective than under uniparental inheritance, irrespective of the size of the bottleneck during uniparental inheritance (Figure 2F and Figure S1). While a tight transmission bottleneck reduces within-gamete variation, the subsequent mixing of cytoplasmic genomes due to biparental inheritance means that cells have higher levels of within-cell variation and lower levels of between-cell variation than under uniparental inheritance (Figure 2A–B).

### 4.4 Varying parameter values does not alter patterns

The choice of fitness function has little effect on our findings (Figure S1). Likewise, varying the selection coefficient does not affect the overall patterns, although the relative advantage of uniparental inheritance over biparental inheritance is larger for higher selection coefficients (Figure S1). Increasing the number of cytoplasmic genomes (*n*) increases the relative advantage of uniparental inheritance over biparental inheritance, whereas increasing the population size (*N*) has little effect (compare Figure S1C with Figure S1A).

### 4.5 Uniparental inheritance helps cytoplasmic genomes purge deleterious substitutions

Free-living asexual genomes accumulate deleterious substitutions more quickly than cytoplasmic genomes when *s*_*FL*_ = 1 (Figure 5A). Biparental inheritance of cytoplasmic genomes causes deleterious substitutions to accumulate more quickly than when inheritance is uniparental (Figure 5). A tight transmission bottleneck slows the accumulation of deleterious substitutions under biparental inheritance, but biparental inheritance always remains less efficient than uniparental inheritance at purging deleterious substitutions (Figure 5).

**Figure 5:**
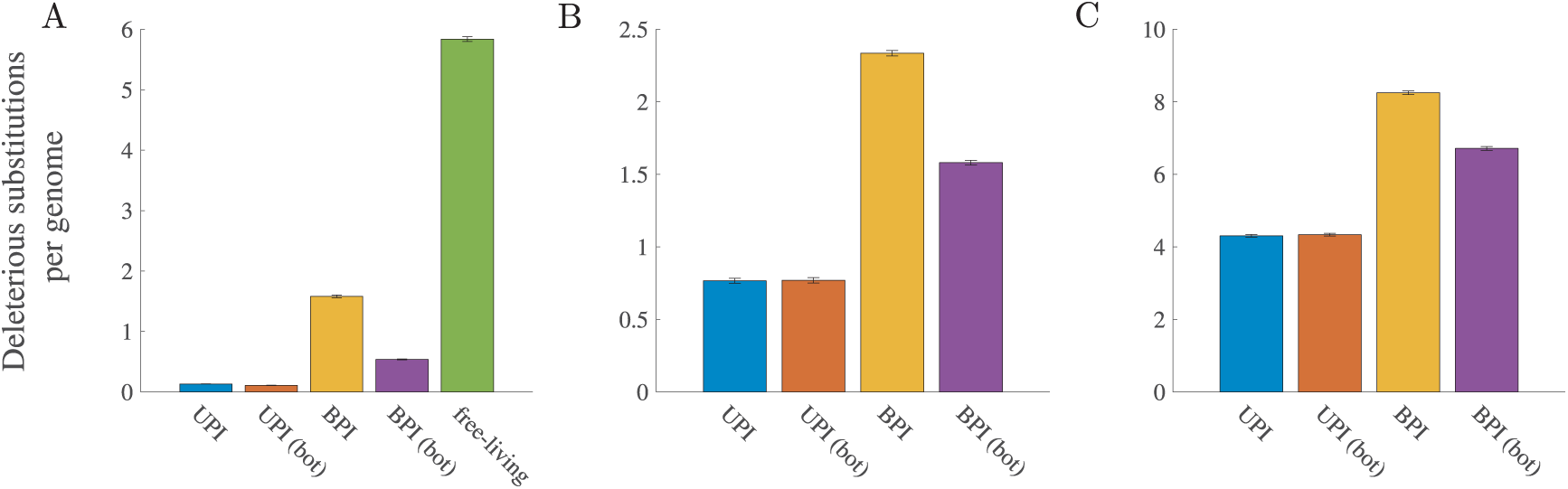
Accumulation of deleterious substitutions in the absence of beneficial mutations. Parameters (unless otherwise stated): *N* = 1000, *n* = 50, *μ* = 10^−7^, a concave down fitness function, and *b* = 25 (relaxed transmission bottleneck) or *b* = 5 (tight transmission bottleneck). UPI: uniparental inheritance with a relaxed bottleneck, UPI (bot): uniparental inheritance with a tight bottleneck, BPI: biparental inheritance with a relaxed bottleneck, and BPI (bot): biparental inheritance with a tight bottleneck. **A**. Comparison with free-living genomes (linear fitness function for both free-living and cytoplasmic genomes, *s*_*d*_ = 0:1, and *s*_*FL*_ = 1). **B**. Mean deleterious substitutions per cytoplasmic genome for *s*_*d*_ = 0:1. **C**. Mean deleterious substitutions per cytoplasmic genome for *s*_*d*_ = 0:01. Error bars are ± standard error of the mean.

### 4.6 Uniparental inheritance reduces hitchhiking of deleterious substitutions

#### 4.6.1 Genetic hitchhiking index

To detect levels of genetic hitchhiking, we developed a method to measure the dependency of deleterious substitutions on beneficial substitutions. When genetic hitchhiking is prevalent, the fixation of deleterious substitutions will more closely follow the fixation of beneficial substitutions relative to random expectation (as the fixation of a beneficial substitution aids the fixation of a deleterious substitution).

We define a “beneficial ratchet” as an event in which the genome that carries the fewest beneficial substitutions is lost from the population. Likewise, we define a “deleterious ratchet” as an event in which the genome carrying the fewest deleterious substitutions is lost. (We describe these events as “ratchets” because a deleterious ratchet is identical to a “click” of Muller’s ratchet (Muller, 1964); a beneficial ratchet is the same concept applied to beneficial substitutions.)

For each simulation, we recorded every generation in which a beneficial ratchet occurred. For each beneficial ratchet, we looked forward in time until we found the nearest deleterious ratchet (including any that occurred in the same generation as a beneficial ratchet). We measured the number of generations separating the beneficial and deleterious ratchet and calculated the mean generations of all such instances.

To obtain a ‘genetic hitchhiking index” (*ϕ*), we divided the mean observed generations separating beneficial and deleterious ratchets by its expectation. The expectation is the mean number of generations that would separate a deleterious ratchet from a beneficial ratchet if deleterious ratchets were randomly distributed through time. If fewer generations separated the beneficial and deleterious ratchets than expected (*ϕ* < 1), we infer that genetic hitchhiking occurred (Figure S2A). If the separation between the beneficial and deleterious ratchets is equal to the expected number of generations (*ϕ* ≈ 1), we infer that beneficial substitutions had no effect on the spread of deleterious substitutions (Figure S2B; see Table S1 for a benchmark of the index). If a greater number of generations than expected separated the beneficial and deleterious ratchets (*ϕ* > 1), we infer that beneficial substitutions inhibited deleterious substitutions (Figure S2C). For details of the genetic hitchhiking index, see Figure S2.

#### 4.6.2 Free-living genomes have higher levels of hitchhiking unless *s*_*FL*_ is high

In all cases, *ϕ* < 1 (Figure 6 and Figure S3), indicating that genetic hitchhiking plays an important role in aiding the spread of deleterious substitutions in both cytoplasmic and free-living genomes. Free-living genomes experience higher levels of hitchhiking than cytoplasmic genomes when *s*_*FL*_ = 1 (Figure 6A). When mutations on free-living genomes have larger effects on fitness, they can experience lower levels of hitchhiking than cytoplasmic genomes under uniparental inheritance (*s*_*FL*_ > 20 in Figure 6B).

**Figure 6:**
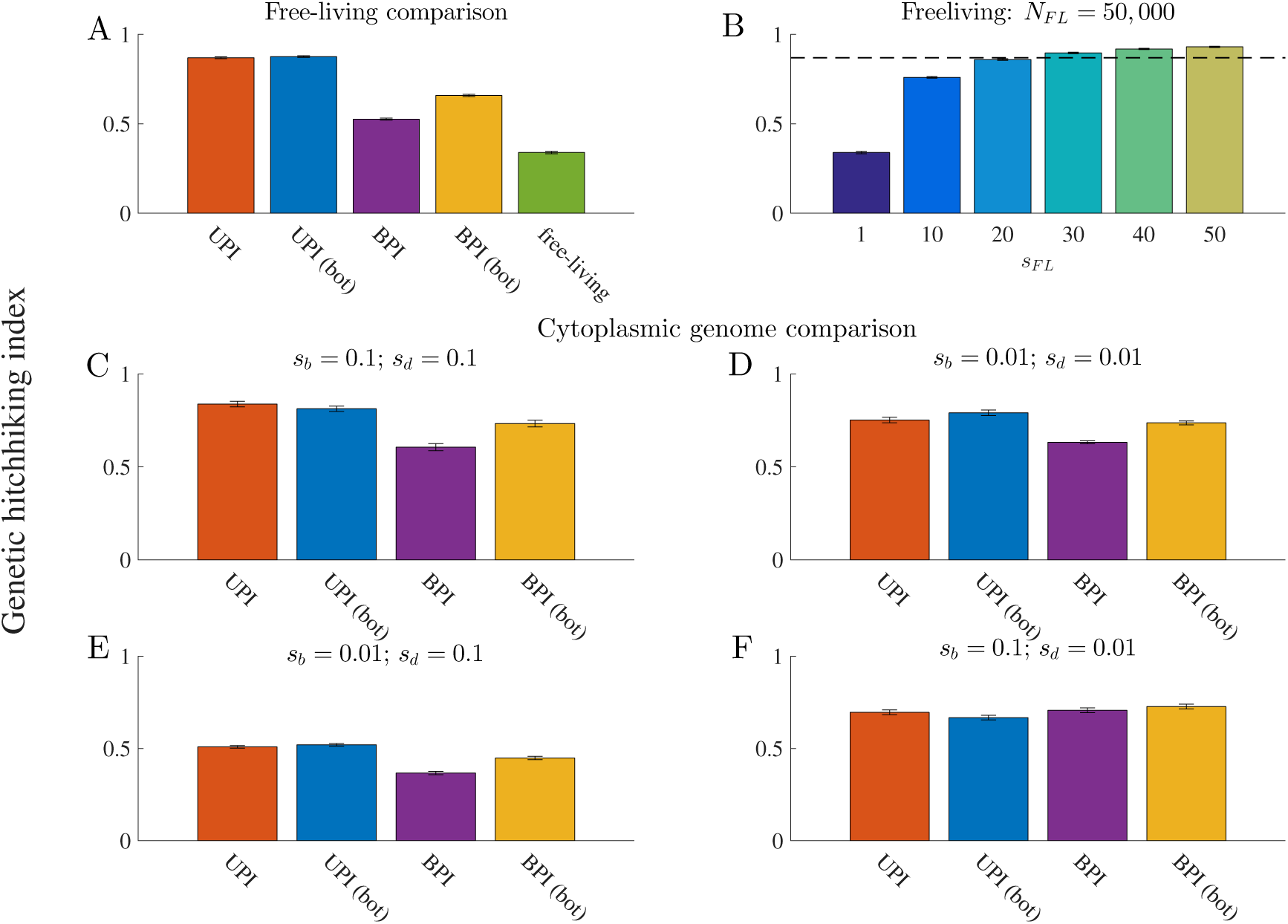
Genetic hitchhiking. The overall level of genetic hitchhiking in each population, measured by our genetic hitchhiking index, *ϕ* (see Figure S2 for details). *ϕ* < 1 indicates the presence of genetic hitchhiking (the lower the value of *ϕ*, the greater the level of hitchhiking). Parameters: *N* = 1000, *n* = 50, μ_*b*_ = 10^−8^, μ_*d*_ = 10^−7^, and *b* = 25 (relaxed transmission bottleneck) or *b* = 5 (tight transmission bottleneck). In all cases, the fitness function for beneficial substitutions is linear. For the free-living comparison in **A–B**, the fitness function for deleterious substitutions is linear, while in the cytoplasmic genome comparison in **C–F**, the fitness function for deleterious substitutions is concave down. UPI: uniparental inheritance with a relaxed bottleneck, UPI (bot): uniparental inheritance with a tight bottleneck, BPI: biparental inheritance with a relaxed bottleneck, and BPI (bot): biparental inheritance with a tight bottleneck. Error bars are ± standard error of the mean. **A**. Free-living comparison, in which *s*_*b*_ = 0:1, *s*_*d*_ = 0:1, *s*_*FL*_ = 1, and *N*_*FL*_ = 50; 000). **B**. Varying the fitness effect of mutations on a free-living genome when *N*_*FL*_ = 50; 000. The dotted line shows the level of hitchhiking for uniparental inheritance (relaxed bottleneck) for comparable conditions (shown in **A**). **C–F**. Genetic hitchhiking in cytoplasmic genomes under different selection coefficients. **C** shows *s*_*b*_ = 0:1 and *s*_*d*_ = 0:1, **D** shows *s*_*b*_ = 0:01 and *s*_*d*_ = 0:01, **E** shows *s*_*b*_ = 0:01 and *s*_*d*_ = 0:1, and **F** shows *s*_*b*_ = 0:1 and *s*_*d*_ = 0:01.

#### 4.6.3 Uniparental inheritance generally reduces levels of hitchhiking

In most scenarios, uniparental inheritance reduces levels of genetic hitchhiking compared to biparental inheritance (Figure 6C–E and Figure S3). The one exception is when *s*_*b*_ > *s*_*d*_, in which case levels of hitchhiking are roughly equivalent under uniparental and biparental inheritance (Figure 6F).

Uniparental inheritance actually increases the proportion of deleterious substitutions that occur concurrently with beneficial substitutions (Figure 7; leftmost bar). This occurs when the genomes that spread carry more than the minimum deleterious substitutions in the population. However, uniparental inheritance also generally increases the proportion of deleterious ratchets in which *ϕ* is large (Figure 7A–C), which occur when the genomes that spread carry the minimum number of deleterious substitutions in the population. Generally, the latter outweigh the former (except for the aforementioned exception), leading to lower levels of genetic hitchhiking under uniparental inheritance (Figure 6 and Figure S3).

**Figure 7:**
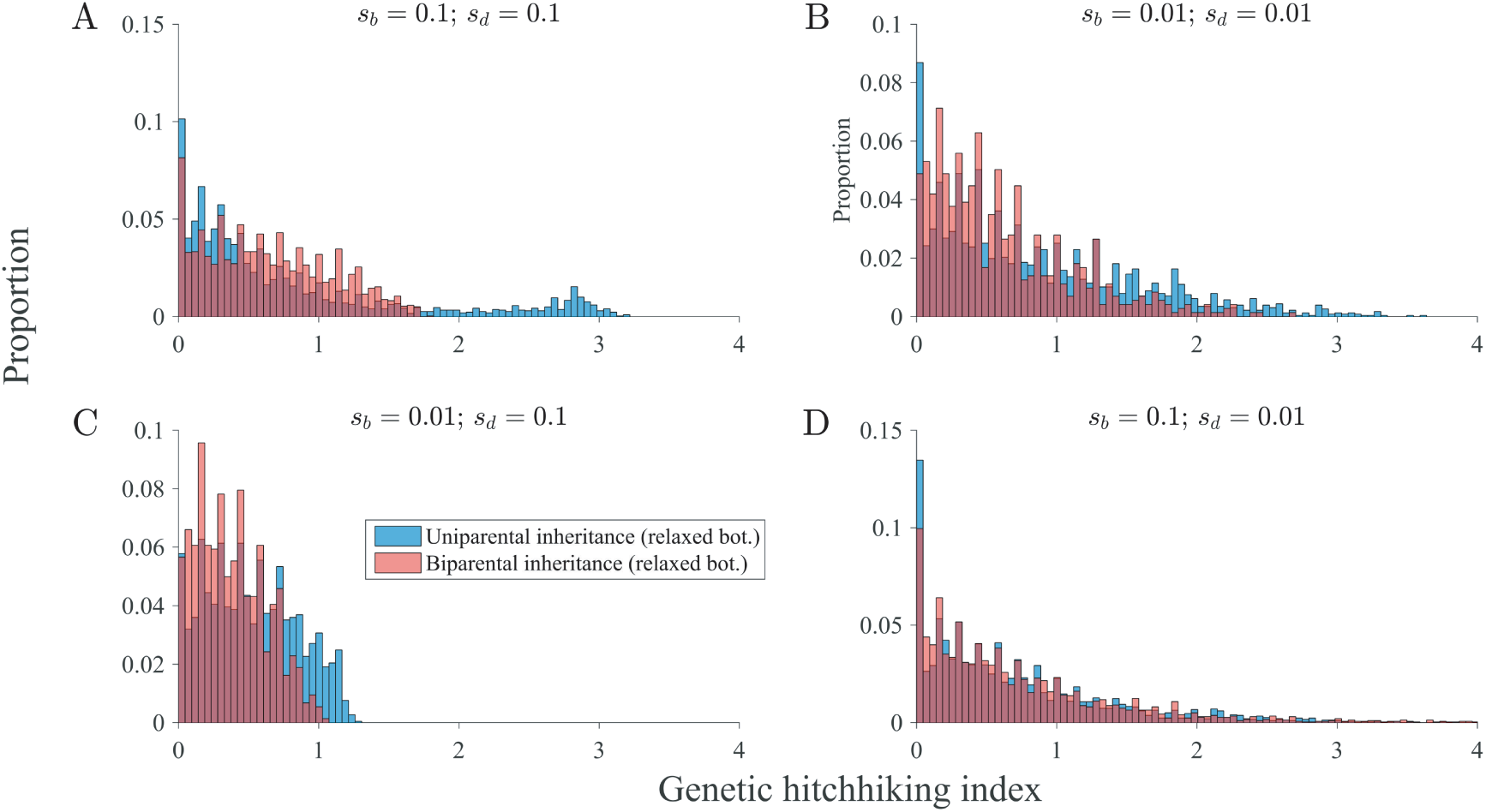
Inheritance mode and the distribution of genetic hitchhiking. The distribution of hitchhiking index values for each pair of beneficial and deleterious ratchets. (A beneficial ratchet occurs when the genome with the fewest beneficial substitutions is lost and a deleterious ratchet occurs when the genome with the fewest deleterious substitutions is lost.) Parameters: *N* = 1000, *n* = 50, μ_*b*_ = 10^−8^, μ_*d*_ = 10^−7^, *b* = 25, a linear fitness function for the accumulation of beneficial substitutions, and a concave down fitness function for the accumulation of deleterious substitutions. **A–D** correspond to the simulations in panels **C–F** in Figure 6. **A**. *s*_*b*_ = 0:1 and *s*_*d*_ = 0:1. **B**. *s*_*b*_ = 0:01 and *s*_*d*_ = 0:01. **C**. *s*_*b*_ = 0:01 and *s*_*d*_ = 0:1. D. *s*_*b*_ = 0:1 and *s*_*d*_ = 0:01. Blue bars pertain to uniparental inheritance, the light pink bars pertain to biparental inheritance, and the dark red bars depict overlapping bars (the dark red bar pertains to whichever color does not show on the top of the bar). We do not plot cases in which the simulation terminates before a beneficial ratchet is followed by a deleterious ratchet. However, we do take these into account when generating the hitchhiking index value: see Figure S2 for details.

### 4.7 Uniparental inheritance promotes adaptive evolution

Cytoplasmic genomes have higher levels of adaptive evolution than free-living genomes unless the effect of mutations on the fitness of free-living cells is much greater than the effect of mutations on eukaryotic host fitness (Figure 8A–C). Among cytoplasmic genomes, uniparental inheritance always leads to higher levels of adaptive evolution than biparental inheritance (Figure 8D–G and Figure S4). While a tight transmission bottleneck combined with biparental inheritance increases the ratio of beneficial to deleterious substitutions, biparental inheritance always has lower levels of adaptive evolution than uniparental inheritance, regardless of the size of the transmission bottleneck (Figure 8D–G and Figure S4).

**Figure 8:**
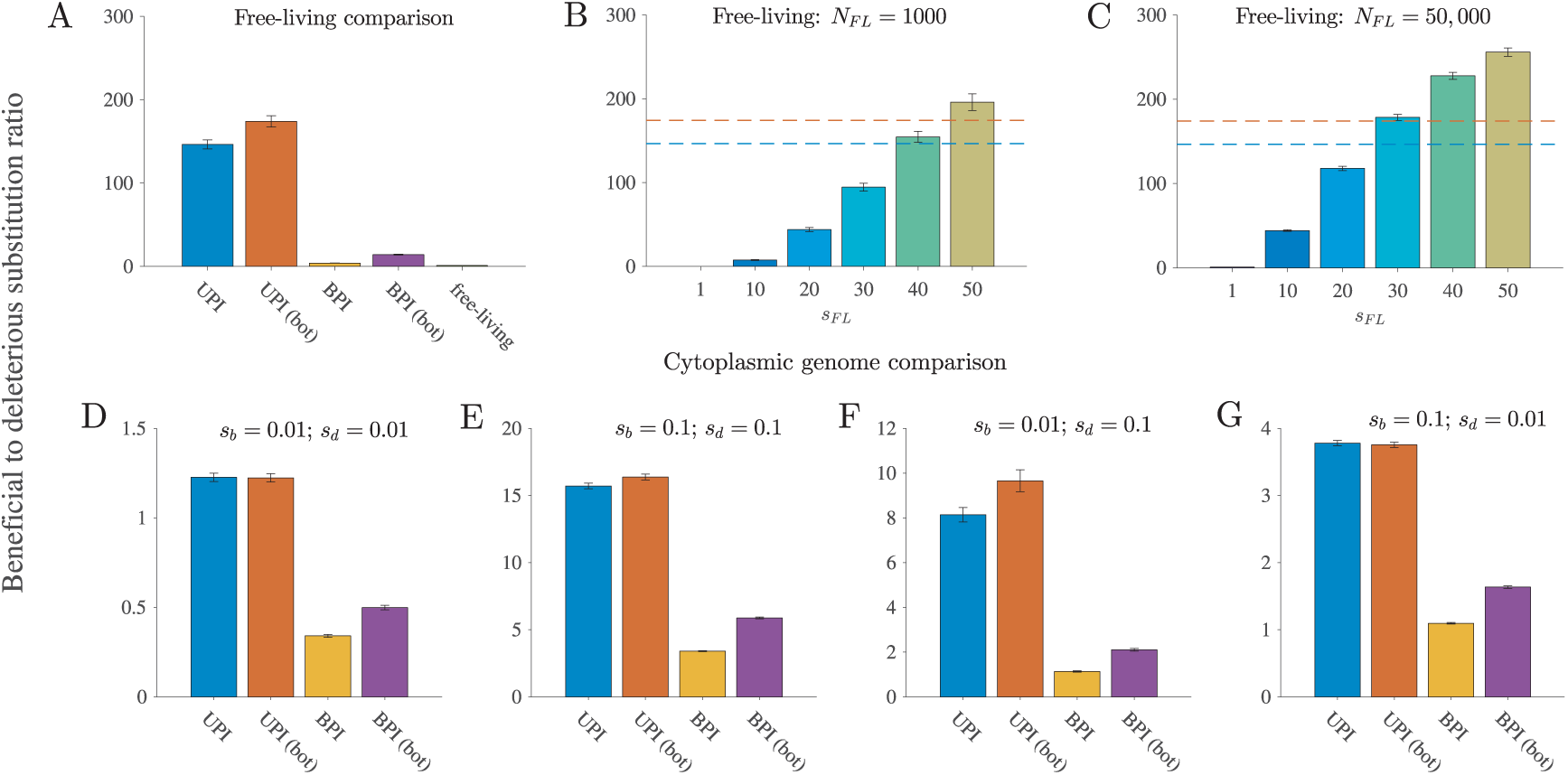
Uniparental inheritance promotes adaptive evolution. Our measure of adaptive evolution is the ratio of beneficial to deleterious substitutions. Parameters (unless otherwise stated): *N* = 1000, *n* = 50, μ_*b*_ = 10^−8^, μ_*d*_ = 10^−7^, *s*_*b*_ = 0:1, *s*_*d*_ = 0:1, and *b* = 25 (relaxed transmission bottleneck) or *b* = 5 (tight transmission bottleneck). UPI: uniparental inheritance with a relaxed bottleneck, UPI (bot): uniparental inheritance with a tight bottleneck, BPI: biparental inheritance with a relaxed bottleneck, and BPI (bot): biparental inheritance with a tight bottleneck. **A**. Comparison with free-living genomes. Here, the fitness function for both beneficial and deleterious substitutions in cytoplasmic genomes is linear. Additional parameters (for free-living genomes only): *N*_*FL*_ = 50; 000, and *s*_*FL*_ = 1. **B–C**. Varying the fitness effect of mutations in free-living genomes relative to cytoplasmic genomes (*s*_*FL*_). The horizontal dotted lines show the ratio of beneficial to deleterious substitutions in UPI (relaxed bottleneck) in blue and UPI (tight bottleneck) in orange depicted in **A**. **B**. Population size of free-living genomes is 1000 (equal to the number of hosts in the UPI and BPI models in **A**). **C**. Population size of free-living genomes is 50,000 (equal to the number of cytoplasmic genomes in the UPI and BPI models in **A**). **D**–**G**. Adaptive evolution in cytoplasmic genomes for a range of selection coefficients. **D**. *s*_*b*_ = 0:01 and *s*_*d*_ = 0:01. **E**. *s*_*b*_ = 0:1 and *s*_*d*_ = 0:1. **F**. *s*_*b*_ = 0:01 and *s*_*d*_ = 0:1. **G**. *s*_*b*_ = 0:1 and *s*_*d*_ = 0:01. To calculate the ratio of beneficial to deleterious substitutions, we first determined the aggregated mean of the number of beneficial and deleterious substitutions for the population at generation 10,000 (average substitutions per cytoplasmic genome). Second, for each of the 500 simulations we divided the mean number of beneficial substitutions per genome by the corresponding mean number of deleterious substitutions per genome. Finally, we took the mean of the ratios of the 500 simulations. Error bars are ± standard error of this mean.

## 5 Discussion

Asexual genomes struggle to accumulate beneficial substitutions and to purge deleterious substitutions (Desai and Fisher, 2007; Felsenstein, 1974; Fisher, 1930; Hill and Robertson, 1966; Lang *et al*., 2013; McDonald *et al*., 2016; Muller, 1932; Park and Krug, 2007). Cytoplasmic genomes, which have a lower effective population size than free-living asexual genomes (Mamirova *et al*., 2007), should be especially susceptible to these limitations of asexual reproduction (Birky, 2008; Lynch, 1996; Rispe and Moran, 2000). These predictions, however, are inconsistent with empirical observations that cytoplasmic genomes can readily accumulate beneficial substitutions and purge deleterious substitutions (Bazin *et al*., 2006; da Fonseca *et al*., 2008; James *et al*., 2016; Popadin *et al*., 2013).

Our study reconciles theory with empirical observations. We show that the specific biology of cytoplasmic genomes increases the efficacy of selection on cytoplasmic genomes relative to free-living genomes when mutations have an equal effect on fitness (i.e. *s*_*FL*_ = 1). By increasing variation in fitness between cells, uniparental inheritance facilitates selection against individuals carrying deleterious substitutions, slowing the progression of Muller’s ratchet. Uniparental inheritance also reduces competition between different beneficial substitutions (clonal interference), causing beneficial substitutions to accumulate on cytoplasmic genomes more quickly than under biparental inheritance.

Uniparental inheritance generally reduces the level of genetic hitchhiking in cytoplasmic genomes, a phenomenon to which asexual genomes are especially susceptible (Lang *et al*., 2013; McDonald *et al*., 2016). Only when beneficial substitutions have a greater effect on fitness than deleterious substitutions does uniparental inheritance not reduce levels of hitchhiking relative to biparental inheritance (Figure 6F). When beneficial mutations have a much stronger effect on fitness than deleterious mutations, it is particularly difficult for asexual genomes to purge deleterious substitutions. Since deleterious substitutions are weakly selected against, they can spread through hitchhiking with beneficial substitutions through positive selection on the latter. Under uniparental inheritance, rapid selective sweeps involving deleterious substitutions may occur too quickly for a new genome—carrying the same number of beneficial substitutions but without excess deleterious substitutions—to be generated and selectively favoured. Nevertheless, of all the genetic hitchhiking scenarios we examined, hitchhiking that involves strongly beneficial and weakly deleterious substitutions is likely the least problematic, as it leads to a net increase in fitness.

By reducing clonal interference, Muller’s ratchet, and in most cases, the level of genetic hitchhiking, uniparental inheritance increases the ratio of beneficial to deleterious substitutions. Both theoretical (Goyal *et al*., 2012) and empirical (Howe and Denver, 2008) evidence suggest that beneficial substitutions can slow Muller’s ratchet by compensating for deleterious substitutions. By increasing the ratio of beneficial to deleterious substitutions, uniparental inheritance effectively increases the ratio of beneficial compensatory substitutions to deleterious substitutions. Thus, the accumulation of beneficial substitutions in cytoplasmic genomes not only aids adaptive evolution (James *et al*., 2016) but improves the ability of cytoplasmic genomes to resist Muller’s ratchet (Bergstrom and Pritchard, 1998; Goyal *et al*., 2012). Together, our findings explain how cytoplasmic genomes are able to undergo adaptive evolution in the absence of sex and recombination.

The effect of a mutation on the fitness of free-living cells (parameter *s*_*FL*_) affects whether adaptive evolution is more efficient in cytoplasmic or free-living genomes. While the comparison between free-living and cytoplasmic genomes helps clarify how the organization of cytoplasmic genomes into hosts affects adaptive evolution, care must be taken when generalizing these findings. First, it is difficult to compare the fitness effects of mutations in free-living and cytoplasmic genomes or to identify a realistic range for *s*_*FL*_. Second, fitness effects of mutations in both free-living and cytoplasmic genomes can differ widely depending on the location of mutations. In mammalian mtDNA, for example, mutations in transfer RNAs (tRNAs) are subject to weaker purifying selection than protein-coding genes (Stewart *et al*., 2008). So while a large s_F L_ value might apply to some mutations, a small *s*_*FL*_ value might apply to others. These variations in fitness effects within animal mtDNA may help explain the different evolutionary trajectories of tRNA and protein-coding genes. While tRNA genes have a substitution rate 5–20 times higher than nuclear DNA (Lynch, 1996), mitochondrial protein-coding genes are more conserved than orthologous genes in free-living bacteria (Mamirova *et al*., 2007) and the genes for nuclear oxidative phosphorylation polypeptides with which they interact (Popadin *et al*., 2013). Ultimately, even when mutations in cytoplasmic genomes have weak effects on fitness, uniparental inheritance will promote adaptive evolution (relative to biparental inheritance) despite these underlying constraints.

We explicitly included a transmission bottleneck as previous theoretical work seemed to suggest that this alone could act to slow the accumulation of deleterious substitutions on cytoplasmic genomes (Bergstrom and Pritchard, 1998). Other work found that host cell divisions—which act similarly to a transmission bottleneck—promoted the fixation of beneficial mutations and slowed the accumulation of deleterious mutations (Takahata and Slatkin, 1983). In contrast, yet another study found that a tight bottleneck increases genetic drift, reducing the fixation probability of a beneficial mutation and increasing the fixation probability of a deleterious mutation (Roze *et al*., 2005). Here we show that these apparently contradictory findings are entirely consistent. We find that a tight transmission bottleneck indeed increases the rate at which beneficial substitutions are lost when rare (Figure 2D). But in a population with recurrent mutation, losing beneficial mutations when rare can be compensated for by a higher rate of regeneration, explaining how a tight bottleneck promotes adaptive evolution despite higher levels of genetic drift. Although a tight transmission bottleneck promoted beneficial substitutions and opposed deleterious substitutions when inheritance was biparental, we show that a bottleneck must be combined with uniparental inheritance to maximize adaptive evolution in cytoplasmic genomes. A transmission bottleneck is less effective in combination with biparental inheritance because the mixing of cytoplasmic genomes after syngamy largely destroys the variation generated between gametes during gametogenesis. For the parameter values we examined, uniparental inheritance is the key factor driving adaptive evolution, as the size of the bottleneck has little effect on the accumulation of beneficial and deleterious substitutions when inheritance is uniparental. It is possible that more extreme transmission bottlenecks (e.g. thousands of genomes down to hundreds or tens) will have a greater effect on adaptive evolution.

We ignored the possibility of within-cell selection between different cytoplasmic genomes. Although within-host replication of cytoplasmic genomes appears to be primarily under host control (Kelly *et al*., 2012; Lee *et al*., 2015), there are several biological examples of “selfish” mitochondrial mutations—those that increase transmissibility of mtDNA but, in doing so, impair host fitness (Clark *et al*., 2012; Gitschlag *et al*., 2016; Ma and O’Farrell, 2016; Taylor *et al*., 2002). Using insights from previous work on two-level selection in cytoplasmic genomes (Rispe and Moran, 2000), we can anticipate how our findings would be affected by within-cell selection. Uniparental inheritance increases variation between hosts and reduces variation within hosts; uniparental inheritance thus increases between-host selection and decreases within-host selection. When within- and between-cell selection act in the *opposite* direction (i.e. fast replicating “selfish” deleterious mutations and slow replicating “altruistic” beneficial mutations (Roze *et al*., 2005)), uniparental inheritance should promote adaptive evolution more efficiently. By minimizing within-cell selection, uniparental inheritance helps prevent mitochondria that carry selfish deleterious mutations from out-competing wild type mitochondria and helps prevent altruistic beneficial mitochondria from being out-competed by wild type mitochondria. When within- and between-cell selection act in the same direction (i.e. “uniformly” deleterious mutations and “uniformly” beneficial mutations (Roze *et al*., 2005)), the outcome is more nuanced. When between-cell selection is much stronger than within-cell selection, uniparental inheritance should promote adaptive evolution. When between-cell selection is much weaker than within-cell selection, however, uniparental inheritance should impair adaptive evolution (relative to biparental inheritance). By minimizing within-cell selection, uniparental inheritance will impede uniformly deleterious mutations from being out-competed by wild type mitochondria and impede uniformly advantageous mutations from out-competing wild type mitochondria.

For simplicity, we ignored recombination in this study. There is an oft-repeated notion in the literature that low levels of recombination, made possible by paternal leakage or occasional biparental inheritance, prevents mitochondrial genomes from accumulating deleterious mutations and succumbing to Muller’s ratchet (Barr *et al*., 2005; Birky, 1995; Greiner *et al*., 2015; Hoekstra, 2000; Neiman and Taylor, 2009). Paternal leakage does occur in animals, and may even be relatively widespread (Dokianakis and Ladoukakis, 2014; Nunes *et al*., 2013; Wolff *et al*., 2013). Recombination between animal mitochondrial DNA has also been observed (Fan *et al*., 2012; Ujvari *et al*., 2007), but it is doubtful whether it is sufficiently frequent to alter evolutionary dynamics (Hagstrom *et al*., 2014; Rokas *et al*., 2003). For example, studies documenting paternal leakage in natural populations have failed to detect recombinant mtDNA (Nunes *et al*., 2013). We have shown that an increase in within-cell variation, which is necessary for recombination among cytoplasmic genomes, reduces the efficacy of selection on hosts and dramatically reduces the level of adaptive evolution in cytoplasmic genomes. Any putative benefits of recombination in alleviating Muller’s ratchet must therefore overcome the acceleration of Muller’s ratchet due to inefficient selection against deleterious mutations. Consequently, we predict that recombination among cytoplasmic genomes will generally hasten Muller’s ratchet rather than slow it.

To our knowledge, the argument that recombination between cytoplasmic genomes can alleviate Muller’s ratchet (Greiner *et al*., 2015; Hoekstra, 2000; Neiman and Taylor, 2009) relies on the findings of models designed for free-living asexual genomes (e.g. (Charlesworth *et al*., 1993; Pamilo *et al*., 1987)) not on models specific to cytoplasmic genomes. This highlights a general finding of our study: population genetic theory developed for free-living genomes cannot be blindly applied to cytoplasmic genomes. Consider effective population size (*N*_*e*_). A lower *N*_*e*_ leads to higher levels of genetic drift (Lynch *et al*., 2006), and it is often assumed that low *N*_*e*_ impairs selection in cytoplasmic genomes (Neiman and Taylor, 2009). However, this assumes that factors which decrease *N*_*e*_ do not alter selective pressures and aid adaptive evolution in other ways. This assumption is easily violated in cytoplasmic genomes, as halving the *N*_*e*_ of cytoplasmic genomes—the difference between biparental and uniparental inheritance—improves the efficacy of selection and can dramatically increase the ratio of beneficial to deleterious substitutions.

The most well-characterized cases of adaptive evolution in cytoplasmic genomes are found in animal mtDNA (Bazin *et al*., 2006; Castoe *et al*., 2008; da Fonseca *et al*., 2008; Grossman *et al*., 2004; James *et al*., 2016; Meiklejohn *et al*., 2007; Ruiz-Pesini *et al*., 2004; Shen *et al*., 2010). For simplicity, our model was based on a single-celled eukaryote life cycle. Multicellular animals, however, differ from single-celled eukaryotes in a number of ways. One difference, in particular, very likely affects adaptive evolution in animal mtDNA. Experiments have shown that pathogenic mtDNA mutations are passed from mother to offspring less frequently than expected by chance, indicating that purifying selection acts within the female germline (Fan *et al*., 2008; Hill *et al*., 2014; Ma *et al*., 2014; Stewart *et al*., 2008). Variation between the mtDNA contents of oocytes, generated by tight bottlenecks during oocyte development, will promote selection between oocytes within the germline (Haig, 2016). Animals may thus be able to select against harmful mtDNA at multiple levels, slowing the progression of Muller’s ratchet.

Although our findings apply most obviously to animal mtDNA, the general insights can be applied broadly to cytoplasmic genomes. In addition to mitochondria, these include plastids and obligate endosymbionts such as *Rickettsia*, *Buchnera*, and *Wolbachia*. Endosymbionts share many traits with cytoplasmic organelles, including uniparental inheritance and multiple copy numbers per host cell. Thus, uniparental inheritance may also be key to explaining known examples of adaptive evolution in endosymbionts (Fares *et al*., 2002; Jiggins, 2006)

## 6 Acknowledgements

We are grateful to Timothy Schaerf for his advice on model design. We thank members of the Behaviour and Genetics of Social Insects Lab and Hanna Kokko for helpful comments on an earlier version of the manuscript. We are also grateful to the anonymous reviewers for thoughtful comments that improved the manuscript. JRC acknowledges funding from the Australian Government (Australian Postgraduate Award), the Society for Experimental Biology, the Society for Mathematical Biology, and the European Society for Mathematical and Theoretical Biology. This work was supported by the Australian Research Council (FT120100120 and DP140100560 to MB) and Intersect Australia (fv4 to JRC and MB). (The support from Intersect Australia was administered through the National Computational Infrastructure (NCI), which is supported by the Australian Government.) We thank The University of Sydney for access to High Performance Computing resources.

## 7 Author contributions

JRC designed the research, performed the experiments, and analyzed the data. JRC and MB wrote the paper.

